# Generation of Heart Organoids Modeling Early Human Cardiac Development Under Defined Conditions

**DOI:** 10.1101/2020.06.25.171611

**Authors:** Yonatan Israeli, Mitchell Gabalski, Kristen Ball, Aaron Wasserman, Jinyun Zou, Guangming Ni, Chao Zhou, Aitor Aguirre

## Abstract

Cardiovascular-related disorders are a significant worldwide health problem. Cardiovascular disease (CVD) is the leading cause of death in developed countries, making up a third of the mortality rate in the US^1^. Congenital heart defects (CHD) affect ∼1% of all live births^2^, making it the most common birth defect in humans. Current technologies provide some insight into how these disorders originate but are limited in their ability to provide a complete overview of disease pathogenesis and progression due to their lack of physiological complexity. There is a pressing need to develop more faithful organ-like platforms recapitulating complex *in vivo* phenotypes to study human development and disease *in vitro*. Here, we report the most faithful *in vitro* organoid model of human cardiovascular development to date using human pluripotent stem cells (hPSCs). Our protocol is highly efficient, scalable, shows high reproducibility and is compatible with high-throughput approaches. Furthermore, our hPSC-based heart organoids (hHOs) showed very high similarity to human fetal hearts, both morphologically and in cell-type complexity. hHOs were differentiated using a two-step manipulation of Wnt signaling using chemical inhibitors and growth factors in completely defined media and culture conditions. Organoids were successfully derived from multiple independent hPSCs lines with very similar efficiency. hHOs started beating at ∼6 days, were mostly spherical and grew up to ∼1 mm in diameter by day 15 of differentiation. hHOs developed sophisticated, interconnected internal chambers and confocal analysis for cardiac markers revealed the presence of all major cardiac lineages, including cardiomyocytes (*TNNT2*^+^), epicardial cells (*WT1*^+^, *TJP*^+^), cardiac fibroblasts (*THY1*^+^, *VIM*^+^), endothelial cells (*PECAM1*^+^), and endocardial cells (*NFATC1*^+^). Morphologically, hHOs developed well-defined epicardial and adjacent myocardial regions and presented a distinct vascular plexus as well as endocardial-lined microchambers. RNA-seq time-course analysis of hHOs, monolayer differentiated iPSCs and fetal human hearts revealed that hHOs recapitulate human fetal heart tissue development better than previously described differentiation protocols^3,4^. hHOs allow higher-order interaction of distinct heart tissues for the first time and display biologically relevant physical and topographical 3D cues that closely resemble the human fetal heart. Our model constitutes a powerful novel tool for discovery and translational studies in human cardiac development and disease.

## INTRODUCTION

CVD and CHD constitute some of the most pressing health problems worldwide due to high prevalence and mortality. Cardiovascular biology researchers and pharmaceutical industries have long relied on transformed cell lines and animal models to study these pathologies; however, the inherent limitations are becoming increasingly clear. In the last decade, and thanks to our increased understanding of developmental pathways and advances in stem cell technologies, the implementation of hPSC-derived cell differentiation protocols has constituted a significant step forward to produce human cells *in vitro* to model human development and disease without the ethical constraints typically associated with human research.

hPSCs enable us to recapitulate some of most important developmental steps *in vitro* to produce specific cardiac cell types with relative ease, high purity and in large amounts^5^. Most of these protocols rely on the controlled stepwise manipulation of the Wnt and BMP signaling pathways using chemical inhibitors^3,6,7^. However, these homogenous cell models are still very far away from the structural and cellular complexity of the tissues and organs they represent (e.g. lack of 3D matrix, disorganized cells, and absence of multicell-type interactions). hPSC-based models of cardiac development and disease frequently disregard the importance of other heart cell types to the disease phenotype, such as the epicardium, cardiac fibroblasts, endothelial cells or endocardial cells. The epicardium significantly contributes to heart metabolism, lipid homeostasis and heart development, and the proepicardial tissue is a main source of cardiac progenitor cells present in multiple cardiac layers^7–9^. Cardiac fibroblasts define the extracellular matrix of the heart and its properties as the heart develops and maturates. Both endothelial and endocardial tissues play fundamental roles in heat homeostasis and blood flow. Remarkable headway has been made in differentiation protocols to produce these specific populations in vitro^7,10–12^, but an integrated approach that aims at cellular complexity and tissue organization is lacking. There is a strong demand to bridge this gap, since producing more faithful *in vitro* models of the human heart will allow us to better model health and disease states for clinical applications.

To circumvent these shortcomings, attempts have been recently made to produce more complex, multicell-type 3D heart tissues or organoids. Tissue engineering approaches, which allow for high control of the end construct, are expensive, work intensive and in most cases not readily scalable or easy to implement. Alternatively, the last decade has seen the emergence of organoid-based approaches prompted by our expanded understanding of human development and the implementation of sophisticated bioengineering methods^13–16^. Organoids are self-assembling 3D cell constructs that recapitulate organ properties and structure to a significant extent (organoid means “resembling an organ”)^16–18^. Organoids have proven particularly useful to study unapproachable disease states in humans or for which animal models are not readily available. While organoids have been used very successfully to model a wide range of organ system (colon, pancreas, kidney)^16,17,19–23^ and disease conditions (cancer, host-microbe interactions)^17,24–28^, their application to cardiovascular studies has been sorely lacking. Cardiac organoid fabrication methods were published as early as 2007^29^, and made noticeable progress in the recent years^30–37^. A variety of organoid fabrication techniques are described in previous research, including the initial 2D differentiation of stem cells to cardiomyocytes followed by dissociation and subsequent aggregation into spheroids^30–32^, which can yield functional organoids, but lack control over the differentiation of non-myocyte cardiac lineages and early development self-organization. The seeding of dissociated stem cells onto a micropatterned platform for further differentiation has been used for precise structural formation of cardiac organoids^29,33–35,^ yet the fallout of this method is the non-physiological physical constraints applied to the developing organoid that may hinder the inherent self-assembling capabilities of differentiating stem cells. Similarly, the manual construction of 3D structures using pre-differentiated cardiomyocytes seeded on a mesh or patch have been recently described^36,37^, which yield 3D tissue constructs that lack the physiological complexity of tissue types and the development of extracellular matrix that the cells form as part of their developmental process. Moreover, these methods can be time-consuming, require significant individual effort and lack cardiac relevant morphology, organization, and cardiac multicell type complexity. Here, we report a novel and optimized hPSC-based heart organoid (hHO) generation protocol with improved cardiac multicell type complexity, higher order structural organization including epicardial and endocardial layers, chamber formation and coronary vascularization. Additionally, our organoid protocol is relatively simple, can be automated, is fully scalable and high-content/high-throughput screening friendly, thus making it readily amenable to pharmacological screening.

## RESULTS

### Differentiation of hPSCs into 3D human heart organoids (hHOs) by Wnt signaling modulation

To develop a protocol for hHO generation using hPSCs we started by using a stepwise chemical modulation of canonical Wnt signaling, an approach inspired by previous monolayer differentiation reports^3,38^. We designed our method with four main goals: 1) high organoid quality and reproducibility; 2) high-throughput/high-content format; 3) relative simplicity (no need for special equipment outside of traditional cell culture instrumentation); 4) defined chemical conditions for maximum control and versatility for downstream applications. We started by assembling hPSCs into spherical aggregates by centrifugation in ultra-low attachment 96-well plates followed by a 48-hour incubation at 37 °C and 95% O_2_ prior to induction. This incubation allowed for spheroid stabilization and was important to increase efficiency, as other incubation times (24 hours, 72 hours) provided inferior results once differentiation started (data not shown). After induction, two-thirds of spent medium was removed and replaced with fresh medium for each medium change, resulting in gradual transitions in exposure to the different signals employed. Optimization of mesoderm and subsequent cardiogenic mesoderm induction was achieved by sequential exposure to two molecules: CHIR99021, a canonical Wnt pathway activator (via specific GSK3 inhibition), and Wnt-C59, a WNT pathway inhibitor (via PORCN inhibition) (**Fig. 1a**). To determine optimal CHIR99021 exposure conditions, we exposed embryoid bodies on day 0 for 24 hours to different concentrations (4 µM, 6.6 µM and 8 µM), followed by Wnt-C59 addition on day 2 for 48 hours. On day 15, hHOs were evaluated for cardiac lineage formation by confocal microscopy using cardiomyocyte specific markers (TNNT2) and image quantification **(Fig. 1b)**. Interestingly, optimal cardiogenic mesoderm induction for all ESC and iPSC lines tested occurred in hHOs that were exposed to lower CHIR99021 concentrations relative to previously reported cardiomyocyte monolayer differentiation protocols, which typically range from 10 to 12 µM CHIR ^3,4,6,7,34,39,40^. A 4 µM CHIR99021 exposure resulted in the highest cardiomyocyte content with 64±5% TNNT2^+^ cells at day 15, versus 9.6±5% and 2.4±2% for 6.6 µM and 8 µM CHIR99021, respectively (**Fig. 1b, d**). This difference is probably due to endogenous morphogen production and paracrine signaling within the developing hHOs, bestowed by the 3D environment and inherent self-assembling properties of the organoids. High magnification immunofluorescence confocal imaging of hHOs stained with TNNT2 antibodies showed clear sarcomere formation and fiber assembly at day 15 (**Fig. 1c**), with most hHOs starting to develop sarcomeres as early as day 7 (**Suppl. Fig 1a)**. Beating hHOs were observed as early as day 6 of the differentiation protocol, with robust and regular beating appearing by day 10 in all samples (**Fig. 1e, f** and **Suppl. Videos 1 & 2**). In general, organoids differentiated from ESCs (H9 cells) exhibited better differentiation efficiencies (**Suppl. Fig. 1b, c)** than iPSC-derived organoids (L1, AICS-37 TNNI1 mEGFP, iPSCORE_16_3). Importantly, we used E8 Flex medium for routine PSC maintenance and embryoid body formation for all successful hHO differentiations, and found that PSCs maintained in mTESR were significantly impaired in forming hHOs using our protocol (radically reduced reproducibility and efficiency)(**Suppl. Fig. 1d-g**). For that reason, all further experiments were performed with hPSCs maintained in Essential 8 Flex medium.

**Figure 1.**
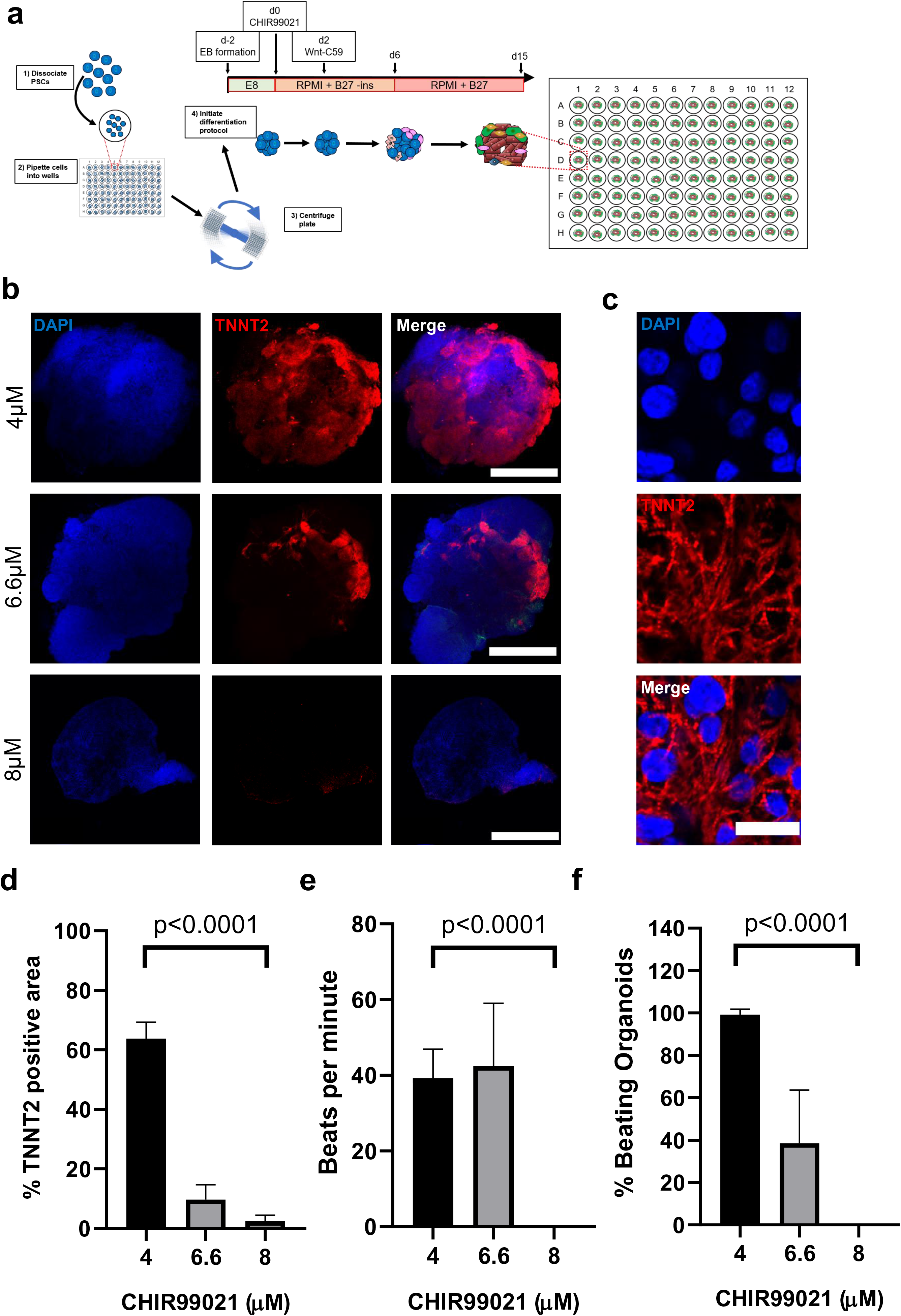
WNT signaling directs cardiomyocyte differentiation in hHOs. **a**, A schematic diagram depicting the protocol used to differentiate TNNT2+ cardiomyocytes in embryoid bodies CHIR99021 concentration is variable at day 0. **b**, Confocal immunofluorescent images for DAPI (blue) and TNNT2 (red), in organoids with CHIR99021 exposure concentrations of 4µM (top), 6.6µM (middle), and 8µM (bottom) at day 15. Scale bars, 500µm. **c**, Confocal immunofluorescent images for DAPI (blue) and TNNT2 (red), in day 15 organoids differentiated using 4µM CHIR showing sarcomere bands (white arrows). Scale bar: 25µm. **d**, Area analysis of cardiomyocyte regions within organoids taken at multiple z-planes as a percentage of TNNT2+ regions to DAPI+ regions of each organoid for the three CHIR99021 concentrations separately. (n=10 for the 4µM CHIR treatment, n=6 for the other two treatments). **e**, Frequency of beats per minute of the hHOs and **f**, percentage of beating hHOs per treatment. (Value = mean ± s.d., two-tailed, unpaired t-test).

### Controlled induction of epicardial lineage in hHOs

To increase organoid complexity and produce more developmentally relevant structures, we decided to adapt methods that have been used successfully in monolayer hPSC differentiation (for specific induction of cardiac lineages other than cardiomyocytes) to our hHO system^41,42^. The method consists of a second activation of canonical Wnt signaling on differentiation days 7-9 to induce secondary cardiac lineages (epicardial cells, cardiac fibroblasts). To determine if this second activation would prime our hHOs to increase complexity and better recapitulate heart development, we tested the effects of a second CHIR99021 exposure on day 7 (**Fig. 2a**). CHIR99021 was added to developing hHOs at varying concentrations (2, 4, 6 and 8 µM, **Fig. 2b**), and exposure lengths (1, 12, 24, and 48 hours, data not shown). Efficiency of epicardial and cardiomyocyte formation was evaluated using confocal imaging and quantification for well-established epicardial (WT1, ALDH1A2, TJP1) and cardiomyocyte (TNNT2) markers at day 15 (**Fig. 2b, c; Suppl. Fig. 2a**). We found that a second CHIR99021 treatment robustly promoted the formation of proepicardium and epicardial cells (**Fig. 2b-d Suppl. Fig. 2a**). Excessively high concentrations or long exposure times led to marked variability in the formation of other cardiac cell types; this variability was especially notable in the cardiomyocytes (**Suppl. Fig. 2b**). We found that hHOs treated with 2 µM CHIR99021 for 1 hour at day 7 had the most physiologically relevant epicardial to myocardial ratio (∼20%), and closest structural layered arrangement found in the heart. This method also produced the most reproducible outcome, with a significant increase in the formation of epicardial cells without affecting cardiomyocyte formation (**Fig. 2b, c; Suppl. Fig. 2c)**. Structurally, epicardial tissue was found predominantly on the surface of the organoids and adjacent to well-defined myocardial tissue areas (TNNT2^+^) (**Fig. 2d**), which recapitulates the structural organization seen in the heart, where the epicardium and myocardium exist as adjacent layers with the epicardium constituting the external layer. The robust expression of TJP1 on epicardial cell membranes also confirmed the epithelial phenotype of these cells (**Fig. 2d**).

**Figure 2.**
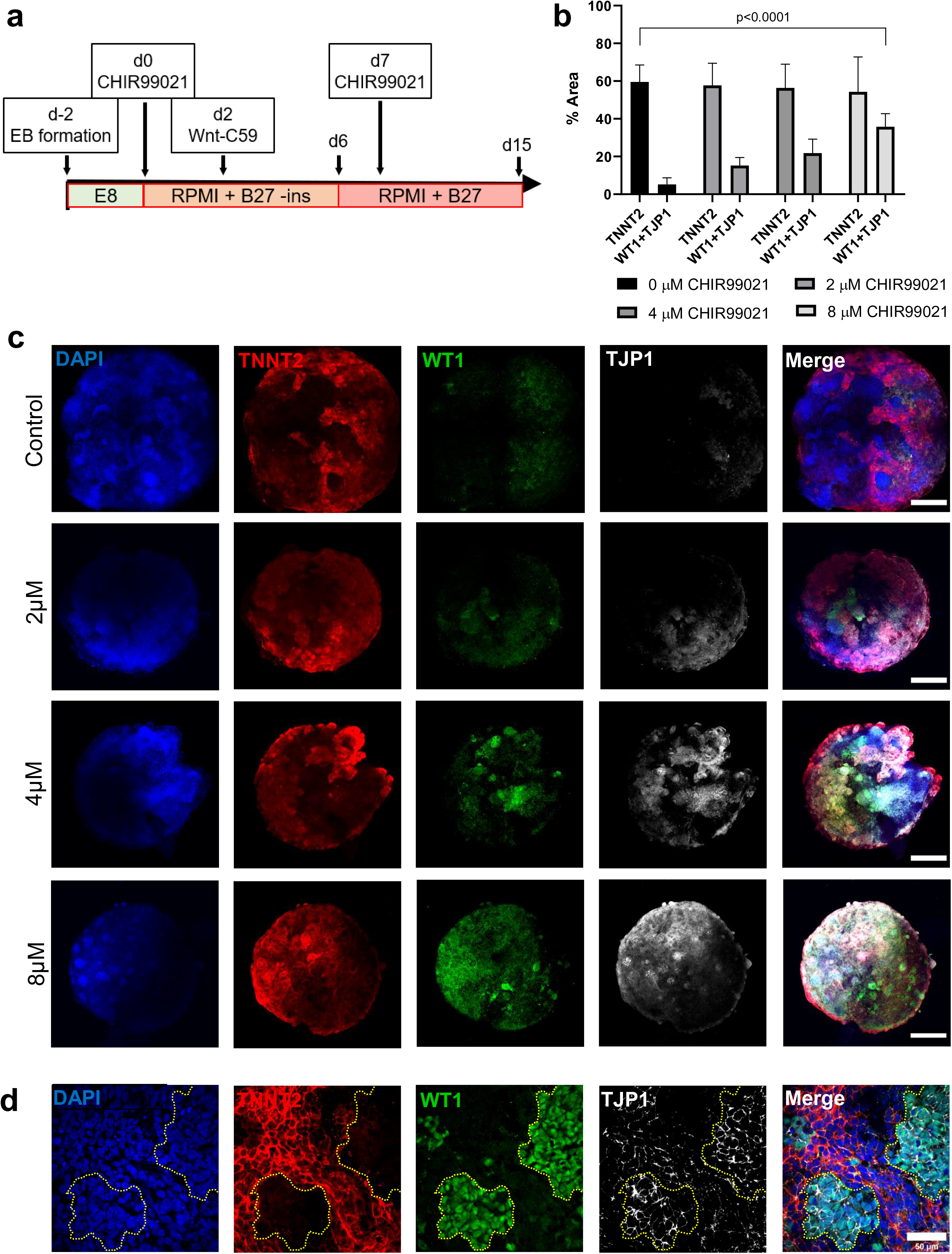
A second CHIR exposure directs epicardial cell differentiation in hHOs. **a**, A schematic diagram depicting the modified protocol used to differentiate TNNT2+ cardiomyocytes and WT1+/TJP1+ epicardial cells in hHOs where x represents the variable CHIR99021 concentration. **b**, Area analysis of cardiomyocyte regions (TNNT2+) and epicardial regions (WT1+ and TJP1+) within organoids taken at multiple z-planes as a percentage of DAPI+ regions of each organoid. (n=7 per condition). **c**, Confocal immunofluorescent images of hHO at differentiation day 15 for DAPI (blue), WT1 (green), TNNT2 (red), and TJP1 (white), with variable concentrations of the second CHIR exposure at day 7 vs control with no second CHIR exposure (scale bars: 500µm), and **d**, high magnification of hHOs with a 2µM second CHIR exposure showing adjacent region of TNNT2+ myocardial tissue and WT1+/TJP1+ epicardial tissue (scale bar: 50µm). (Value = mean ± s.d., two-tailed, unpaired t-test).

### Transcriptomic analysis reveals hHOs closely model human fetal cardiac development and produce all main cardiac cell lineages

To characterize hHOs at the cellular and molecular level, we performed detailed RNA-sequencing (RNA-seq) analysis at different stages of organoid formation and development. hHOs were collected at different timepoints (through day 0 to day 19) of differentiation (**Fig. 3**). Unsupervised K-means cluster analysis revealed organoids progressed through three main developmental stages: day 0 - day 1, associated with pluripotency and early mesoderm commitment; day 3 – day 7, associated with early cardiac development; and day 9 – day 19, associated with fetal heart maturation (**Fig. 3a, Suppl. Fig. 3**). Gene ontology biological process analysis identified important genetic circuitry driving cardiovascular development and heart formation (**Fig. 3a**; **Suppl. Data Table 1**, raw data deposited in GEO under GSE153185). To compare cardiac development in hHOs to that of existing methods, we performed RNA-seq on monolayer iPSC-derived cardiac differentiating cells using well-established protocols^3^, along with using publicly-available data from previous reports using different monolayer cardiac differentiation protocols and also fetal heart tissue (gestational age days 57-67)^43^ (GSE106690). In all instances, activation of hHO cardiac development transcription factor expression regulating first and second heart field specification (FHF, SHF, respectively) was similar to that observed before in monolayer PSC-derived cardiac differentiation, corresponding well to that observed in fetal heart tissue (**Fig. 3b**). Interestingly, gene expression profiles showed hHOs had a higher cardiac lineage cellular complexity than cells that underwent monolayer differentiation, including epicardial, endothelial, endocardial, and cardiac fibroblast populations (**Fig. 3c**). These data suggest a significant enrichment in the structural and cellular complexity of our hHOs, thus bringing them in line with fetal hearts. This was confirmed by extending our gene expression analysis to look at several widespread critical gene clusters involved in classic cardiac function (**Fig. 3d**), including conductance, contractile function, calcium handling and cardiac metabolism among others. Of special interest, hHOs produced significant amounts of heart-specific extracellular matrix, a feature that was present in the fetal hearts but totally absent in monolayer differentiation protocols (**Fig. 3d**). Overall, hHOs had individual expression profiles better matching those of fetal hearts, and the global hHO transcriptome was closer to that observed in fetal hearts than any of the monolayer protocols achieved, as determined by hierarchical clustering (**Fig. 3e)**.

**Figure 3.**
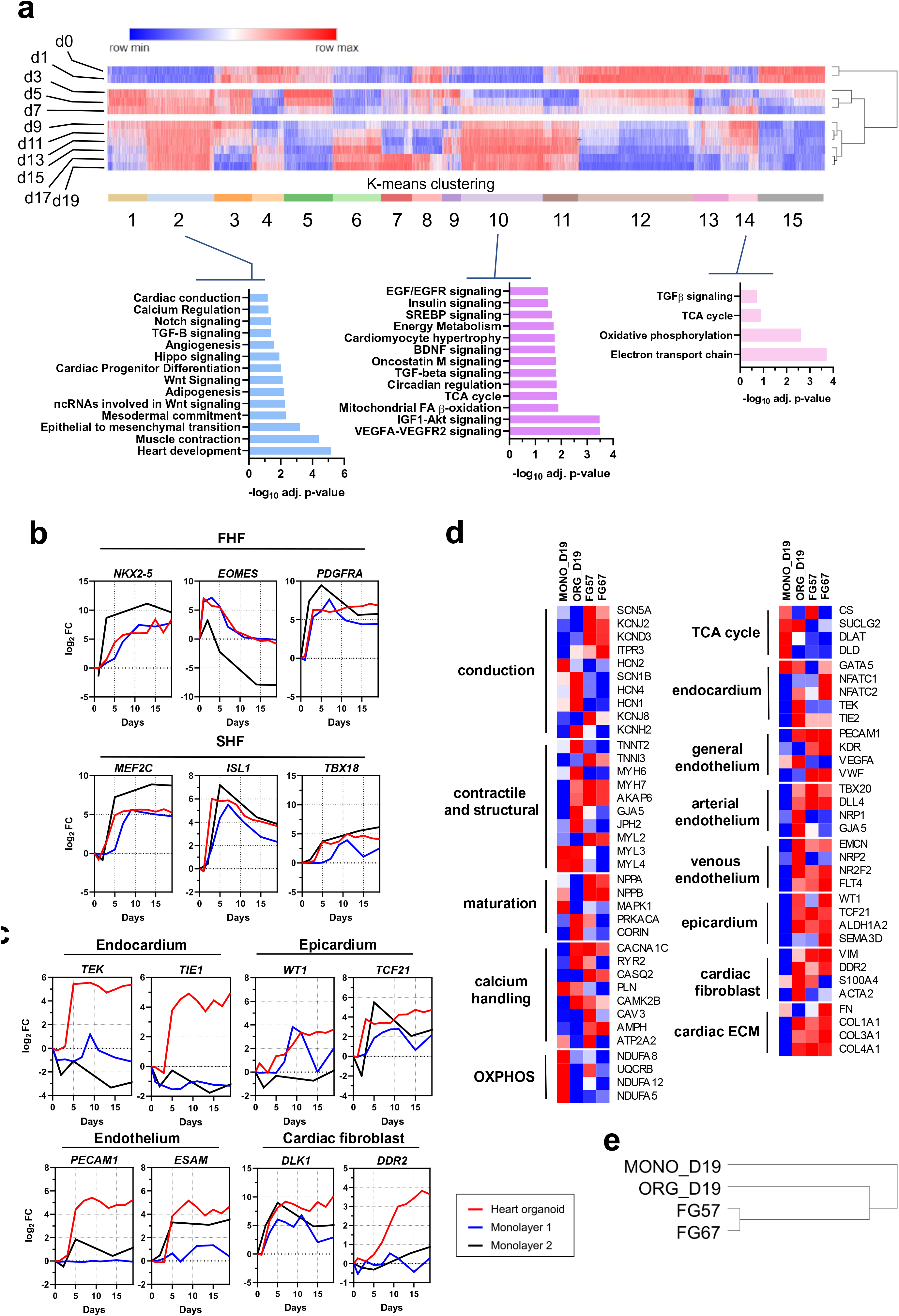
Transcriptomic analysis indicates heart organoids recapitulate multicell type complexity, development and maturation steps similar to embryonic fetal hearts. **a**, K-means cluster analysis of heart organoid transcriptomes by RNA-seq. Clusters strongly associated with fetal heart development (e.g. 2, 10 and 14) appear from day 9 onwards. Pathway enrichment analysis is provided below for representative cardiac-specific clusters. **b**, Gene expression analysis (log_2_ fold-change vs. D0) of first and second heart field markers over heart organoid differentiation process (FHF, SHF respectively). **c**, Gene expression analysis (log_2_ fold-change vs. D0) for cardiac-specific cell type populations in heart organoids, including epicardial cells, fibroblasts, endocardial cells and endothelium. **d**, Normalized comparison of key genes involved in cardiac function across heart organoids, monolayer differentiation methods and fetal hearts at gestational day 57-67. **e**, Hierarchical clustering analysis of heart organoids, monolayer differentiation and fetal hearts.

### A second Wnt activation leads to the formation of multicell type cardiac lineages and emergence of cardiac fetal-like morphological and structural 3D complexity

The results from transcriptomic analysis suggested the second CHIR99021 exposure led to the formation of other mesenchymal derived lineages and higher complexity in hHOs. To better evaluate this finding, we performed immunofluorescence analysis for additional cardiac cell lineages. Confocal imaging revealed a robust interconnected network of endothelial cells (PECAM1^+^), and vessel-like formation throughout the organoid (**Fig. 4a**). Higher magnification uncovered a complex web of endothelial cells adjacent or embedded into myocardial tissue (**Fig. 4b**). 3D reconstruction of confocal imaging stacks showed a well-connected endothelial network intertwined in the hHO tissue (**Fig. 4c, Supp. Videos 3-5**). These results strongly indicate that during hHO development, self-organizing endothelial vascular networks emerge in response to the 3D cardiovascular environment, adding a coronary-like vascular network to the organoids, a phenomenon not observed before. We found the vessels formed by PECAM1+ cells had very consistent diameters of 9.5±0.99 µm and 9.1±0.65 µm (for iPSC-L1 and ESC-H9 respectively), sufficient for functional blood flow (**Suppl Fig. 4a, Suppl. Videos 4**,**5)**. Confocal microscopy revealed well-developed lumens for blood vessels throughout the hHOs (**Fig. 4a)**. Further confocal imaging confirmed the presence of cardiac fibroblasts positive for THY1 and VIM (**Fig. 4d**), which made up 12±2% of the tissues in the hHOs (**Fig4 c,e**), similar to the composition of the fetal heart described in the literature^44^. We suspected that the cavity-like spaces observed within the TNNT2+ tissue might possess chamber-like qualities and mimic early heart chamber formation. Immunofluorescence analysis of the endocardial marker NFATC1 revealed the formation an endocardial layer of NFATC1^+^ cells lining the chambers embedded in the TNNT2^+^ tissue, similar to the endocardial lining of the heart (**Fig. 4e, Suppl. Fig 4d**). Analysis of hHO development revealed the formation of chambers as early as day 5 of the differentiation protocol, with a continuous increase in both chamber size in proportion of hHO area through day 15 (**Fig. 5a)**. We employed optical coherence tomography (OCT) to characterize chamber properties using minimally invasive means, thus preserving chamber physical and morphological properties. OCT showed clear chamber spaces within the hHOs, typically with one or two larger chambers near the center of the organoids, and several smaller ones near the edges (**Fig. 5b, Suppl. Fig**.**5**). 3D reconstruction of the internal hHO topology revealed a high degree of interconnectivity between chambers (**Videos 6-8**). Expression of cardiac-specific ECM genes in the hHOs resembling the fetal heart matrix, such as *COL1A1, COL4A1, COL5A2, FBN1, EMILIN1, HSPG2* and *LAMA2* might be an important factor in chamber organization and deserves further examination in the future (**Fig 5c)**.

**Figure 4.**
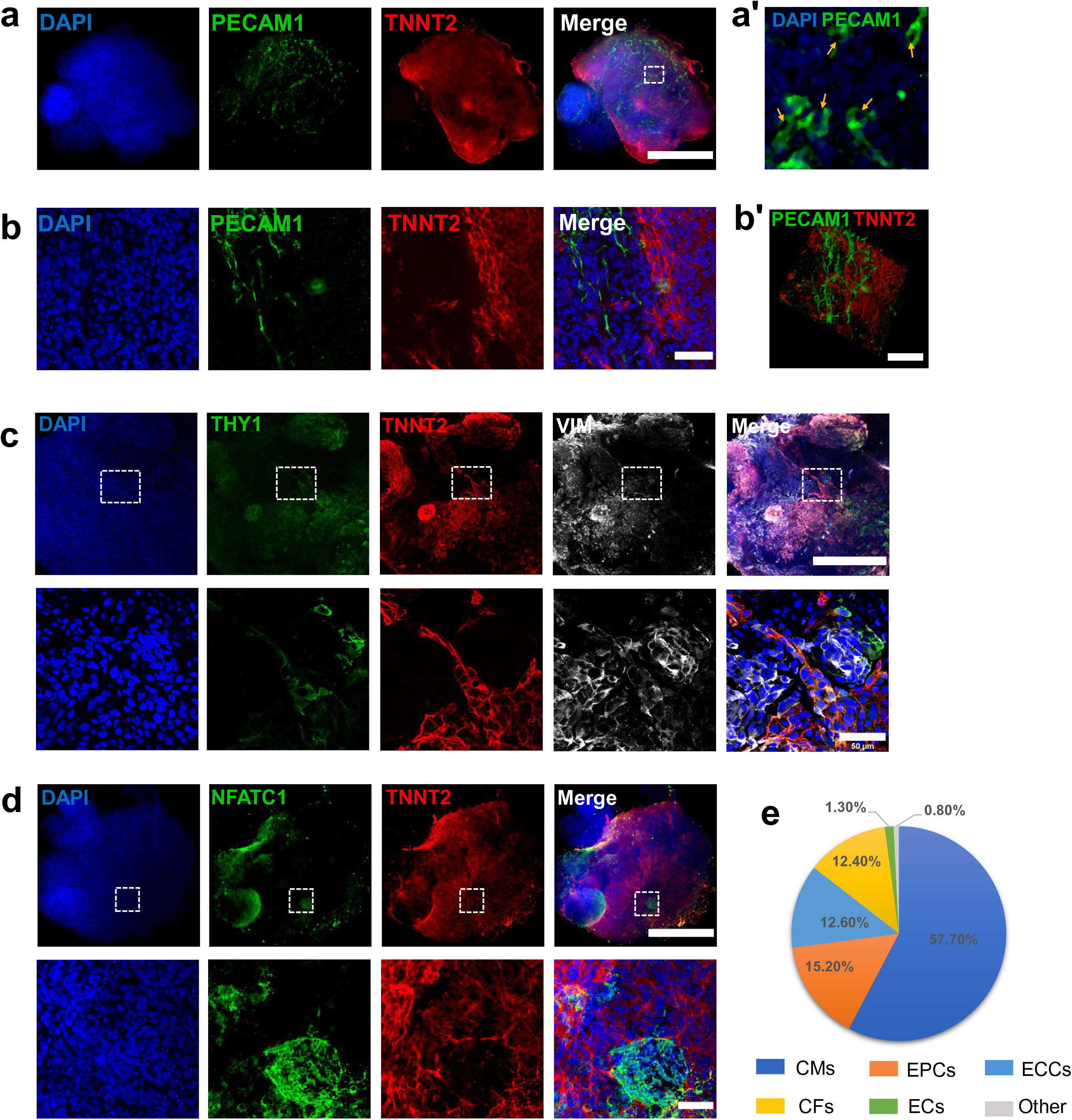
hHO cardiac cell lineage composition. **a-e**, immunofluorescence images of various cell lineages composing the hHOs. **a**, Endothelial marker PECAM1 (green) showing a defined network of vessels throughout the organoid and adjacent to TNNT2+ (red) tissue, DAPI (blue); Scale bar: 500µm. **b**, 60X magnification of PECAM1+ endothelial tissue in close proximity to TNNT2+ myocardial tissue (scale bar: 50µm) **c**, Cardiac fibroblast markers THY1 (green) and VIMENTIN (white) present throughout the hHOs, TNNT2+ (red), DAPI (blue); Scale bar: 500µm, inset: 50µm. **d** endocardial marker NFATC1 (green) highly expressed within microchambers of TNNT2+ (red) tissue (scale bar: 500µm, inset: 50µm). **e**, Pie chart of average cell composition in hHOs, calculated as the percentage of whole organoid area using ImageJ.

**Figure 5.**
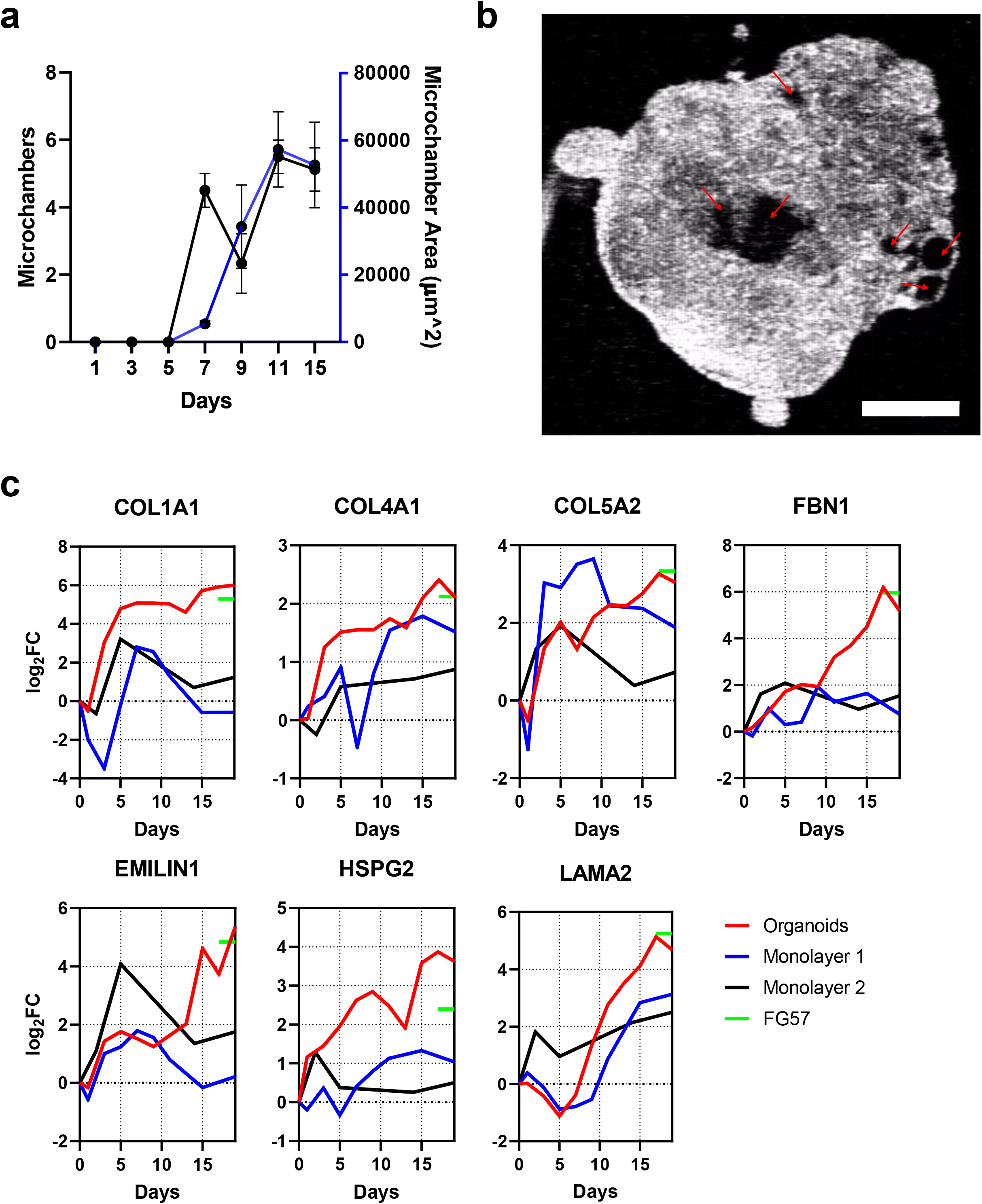
Microchamber formation and ECM proteins. **a**, Number and size of microchambers (left and right axes, respectively) between days 0 and 15 of differentiation protocol. **b**, Optical coherence tomography images showing a cross-section of the center of the organoid, revealing microchambers (red arrows, scale bar: 250µm). **c**, Gene expression analysis (log_2_ fold-change vs. D0) of ECM proteins in hHOs (red) compared to monolayers (black and blue) and fetal heart expression at gestational day 57 (green).

### BMP4 and activin A improve heart organoid chamber formation and vascularization

The growth factors bone morphogenetic protein 4 (BMP4) and activin A have frequently been used as alternatives to small molecule Wnt signaling manipulation since they are the endogenous morphogens that pattern the early embryonic cardiogenic mesoderm and determine heart field specification in vivo^27,45–47^. We wondered whether BMP4 and activin A, in combination with our small molecule Wnt activation/inhibition protocol, could synergistically improve the ability of hHOs to recapitulate cardiac development in vitro. We tested the effect of BMP4 and activin A in the context of our optimized protocol by adding the two morphogens at 1.25 ng/ml and 1 ng/ml, respectively (recommended concentrations found in the literature^27^), at differentiation day 0 in conjunction with 4 µM CHIR99021. No significant differences were found in formation myocardial (TNNT2*+)* or epicardial (WT1+/TJP1*+)* tissue between control hHOs and treated ones (**Fig. 6a**). However, significant differences in organoid size (hHOs treated with growth factors were about 15% larger in diameter) **(Fig. 6b, c)**, which may correspond with the increase in microchamber size and connectivity(BMP4/Activin A were ∼50% more interconnected with other chambers (**Fig. 6d, e, g)**, compared to control hHOs)., were observed. Confocal analysis by immunofluorescence of hHOs treated with BMP4 and activin A showed a 400% increase in the area of PECAM1+ tissue, indicating a significant effect on organoid vascularization (**Fig. 6f, h**), which might account for the increase in hHO size by facilitating gas diffusion and transport of nutrients.

**Figure 6.**
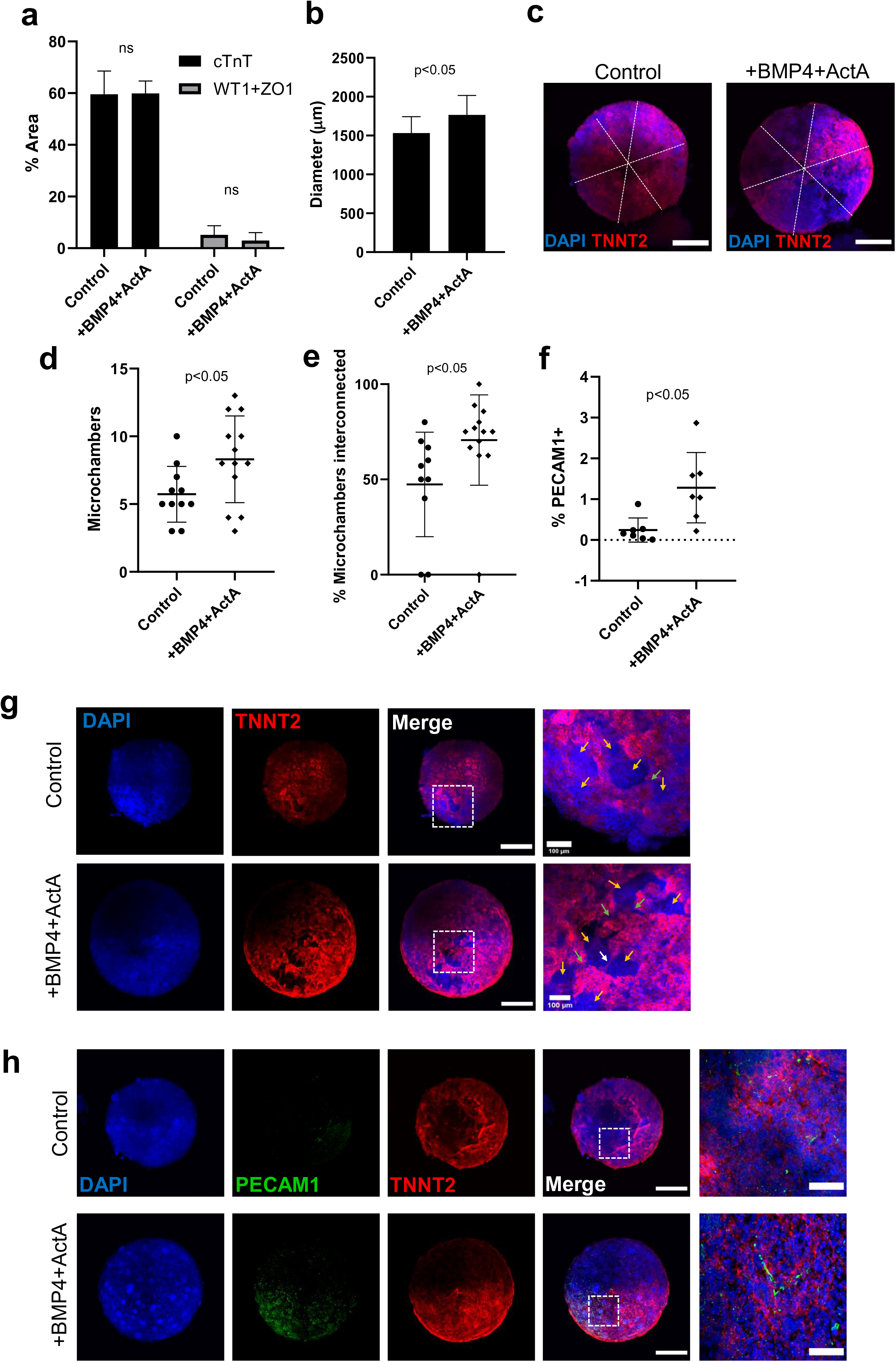
BMP4 and Activin A affect cardiac differentiation and development. **a-j**, all panels compare hHOs differentiated with CHIR alone (Control) and with CHIR+BMP4+Activin A (Treated). **a**, Area of cardiomyocyte and epicardial positive regions as a percentage of total organoid area and **b**, organoid diameter, (*n*=8 per condition). **c**, dashed lines showing the diameter of a control (left) and treated (right) organoid averaged to determine the diameter. **d-e**, *n*=12 per condition. **d**, Number of microchambers in TNNT2+ areas, and **e**, interconnectivity of microchambers measured by separation of microchambers by thin TNNT2+ filaments or by thin channels showing clear connection. **f**, PECAM1+ tissue as a percentage of total organoid area, (measured using MaxEntropy threshold on ImageJ and analyzed all particles of 25µm^2^ to avoid small speckles, *n*=7 per condition). **g**, Immunofluorescence images of hHOs showing interconnected microchambers (yellow arrows), *TNNT2*+ filaments (white arrows), and channels connecting microchambers (green arrows), DAPI (blue), *TNNT2* (red), scale bar: 500µm, inset: 100µm **i**, Immunofluorescence images of hHOs showing DAPI (blue), PECAM1+ tissue (green), and *TNNT2*+ tissue, scale bar: 500µm, inset: 50µm. (Value = mean ± s.d., two-tailed, unpaired t-test).

## Discussion

In recent years, hPSC-derived cardiomyocytes have become critically useful tools to model aspects of heart development^39,48,49^, human genetic cardiac disease^50,51^, therapeutic screening^32,52^, and cardiotoxicity testing^53–55^. Nonetheless, the complex structural morphology and multitude of tissue types present in the heart impose severe limitations to current *in vitro* models. Here we sought to create a highly reproducible, scalable, novel differentiation protocol, to create physiologically relevant human heart organoids with high structural and multicell type complexity using hPSCs. We optimized defined PSC culture conditions and created and optimized a multistep manipulation conditions for canonical Wnt signaling using GSK3 and *PORCN* inhibitors. These conditions lead to the formation of most cardiac lineages in a self-assembling heart organoid with similar properties to the fetal heart. This method consistently yields cardiac organoids comprised of about 59% cardiomyocytes, 15% epicardial cells, 13% endocardial cells, 12% cardiac fibroblasts and 1% endothelial cells, by area, and show robust beating throughout the entire hHO within a week from differentiation initiation. Interestingly, hPSCs cultured in mTeSR rather than E8 culture medium prior to differentiation yielded significantly less reproducible hHOs, less control over cardiac lineages and poor beating, suggesting components of mTESR, such as albumin, might be interfering with the differentiation protocol. hHOs were successfully derived from three independent iPSC lines and one ESC line, demonstrating reproducibility. The fetal-like morphology of the cardiomyocytes and the self-assembling nature of the hHOs allude to a complex three-dimensional structure containing a multitude of cardiac cell lineages allowing for higher order interactions between different heart tissues. When compared with existing cardiomyocyte monolayer differentiation methods, hHOs showed higher expression of genes associated with conduction, contractile function, calcium handling, and various cardiac cell populations, which better resembles gene expression data retrieved from human fetal hearts. The depiction of a complex transcriptome highly recapitulative of human fetal heart tissue further strengthens the complexity and validity of the hHO as a model of human heart development.

The epicardium, an epithelial layer that encapsulates the human heart, is involved in many important heart processes, including heart development, metabolism, lipid homeostasis and myocardial injury responses^8,56^. Epicardial signaling cascades are essential for cardiac lineage specification^8^. During embryonic development, cells from the proepicardial organ (PEO), a cluster of embryonic cells^56^, migrate to the surface of the heart to form the epicardium, and some of these cells can undergo EMT to generate other cardiac lineages including cardiac fibroblasts^8,56,57^. Due to its capacity to communicate with the myocardium and its ability to mobilize stem cell populations, the epicardium has become a key focus of research in cardiac regeneration and repair^8,42,56,58,59^. The epicardium also plays a fundamental but underexplored role in multiple types of cardiovascular and metabolic disease, including diabetic cardiomyopathy, coronary artery disease and metabolic syndrome. In this last condition, epicardial-derived fat experiences a significant expansion and correlates strongly with morbidity, highlighting the potential relevance of the epicardium to these conditions. Inspired by a previous epicardial differentiation method^42^ based on a second Wnt activation step at day 7 of differentiation, we created and optimized conditions for producing heart organoids with well-defined regions of epicardial tissue adjacent to myocardial tissue. This advancement will facilitate the study and modeling of physiologically relevant epicardial-myocardial interactions *in vitro* relevant for health and disease.

An acute limitation of many organoid systems is a lack of a functional vascular network to facilitate the exchange of nutrients and removal of waste material and instead rely solely on diffusion^60–62^. Several vascularized organoids have been described in the literature including brain^61^, kidney^63^, and blood vessel^10^, however none to this point with cardiac organoids. In these studies, various techniques are used to induce vascularization including implantations in mice^61^, culturing the organoids under flow^63^, and embedding endothelial cells in a Matrigel/collagen matrix and inducing their migration^10^ to create a vascular network. Remarkably, we observed the formation of a robust interconnected vascular plexus in our final protocol for hHOs without any additional steps. Further studies into the functionality of this vascular tissue will be necessary, particularly to determine the degree of maturation of these vessels, and if they closely resemble coronary vasculature -- a feature that would open the door to modeling coronary vasculature pathologies that arise due to cardiovascular disease and metabolic disorders. In addition to vasculature, we also observed the spontaneous reorganization into interconnected chambers, a very powerful 3D feature indicative of recapitulation of fetal-like organogenesis. Previous studies into microchamber formation *in vitro* utilized micropatterning of hPSCs into a confined area to generate 3D cardiac microchambers with cell-free regions, a myofibroblast perimeter and nascent trabeculae^33^. While the previously defined spatial patterning of the cardiac organoids showed some fetal-like formation of cardiac microchambers, they lacked endocardial tissue^34^; a crucial playmaker in heart maturation and morphogenesis^64^. The hHOs reported here form multiple microchambers which are lined with NFATC1+ endocardial cells and are interconnected as seen in the OCT cross-sectional imaging (**Suppl. Videos 6-8**). This fact, coupled with the physically unconstrained development of the organoids, reveals a more fetal-like maturation of the heart tissue, supported by our gene expression data **(Fig. 3)**.

The important role that cardiac fibroblasts play on cardiac development and cardiac matrix production/organization is often overlooked in *in vitro* models. Most cardiac fibroblasts in embryonic development arise from the PEO^57,65,66^, highlighting the necessity of proepicardial induction in developmental heart models. Immunofluorescence analysis of our hHOs revealed the presence of cardiac fibroblast markers including the membrane glycoprotein Thy1, which is involved in cell-cell and cell-matrix adhesion^66,67^, and the intermediate filament protein Vimentin, typically seen in cells of mesenchymal lineage^67^. Other cardiac fibroblast markers were found in the hHOs via RNA-sequencing analysis, including DDR2^66^ which plays an important role in EMT and the FHF marker PDGFRα which is crucial for vascularization during development and is also a strong marker of cardiac fibroblasts^67^. All these provide strong indication of the increased complexity of out hHO system and the closer resemblance to fetal heart tissue.

Together with the use of small molecule inhibitors that manipulate canonical Wnt signaling, successful cardiomyocyte differentiation has been achieved in the past using morphogens such a BMP4 and activin A^3,39^, which lead to the induction of cardiac mesoderm in the embryo^68^. Established differentiation protocols using these growth factors show effective differentiation to various cardiac mesoderm progenitors^4,68,69^. Recently, gradient exposures to specific concentrations of BMP4 and activin A have been explored in the specification of first and second heart field formation^27^. The addition these growth factors to the initial CHIR exposure in our hHO differentiation protocol led to improved morphological features such as increased microchamber interconnectivity and vascularization.

In summary, we describe here a highly reproducible and high-throughput human heart organoid derivation method, with multicell type and morphological complexity closely recapitulating the developing human fetal heart. This model constitutes a valuable tool to investigate the development of the human heart and the etiology of congenital heart defects. Furthermore, refinement and improved maturation protocols might allow to model adult cardiac settings, such as cardiotoxicity screening and cardiovascular-related disorders.

## Acknowledgments

We wish to thank the MSU Advanced Microscopy Core and Dr. Jackson at the Department of Pharmacology and Toxicology for access to confocal microscopes, and the MSU Genomics Core for sequencing services. We also wish to thank all members of the Aguirre Lab for valuable comments and advice.

## Author contributions

Yonatan Israeli and Aitor Aguirre designed all experiments and conceptualized the work. Yonatan Israeli performed all experiments and data analysis. Mitchell Gabalski and Aaron Wasserman performed PSC and organoid culture. Kristen Ball performed work on epicardial differentiation optimization in hHOs. Jinyun Zou, Guangming Ni and Chao Zhou performed optical coherence tomography experiments and data analysis. Aitor Aguirre supervised all work.

## Sources of Funding

Work in Dr Aguirre’s laboratory was supported by the National Heart, Lung, and Blood Institute of the National Institutes of Health under award number HL135464 and by the American Heart Association under award number 19IPLOI34660342. Work in Dr. Zhou’s laboratory was supported by grants from the National Institutes of Health under award number R01EB025209.

## Disclosures

The authors declare no conflicts of interest.

## Materials and Methods

### Stem cell culture

Human iPSC lines used in this study were iPSC-L1, AICS-0037-172 (Coriell Institute for Medical Research, alias AICS), iPSCORE_16_3 (WiCell, alias iPSC-16) and human ESC line H9. All PSC lines were validated for pluripotency and genomic integrity. hPSCs were cultured in Essential 8 Flex medium containing 1% penicillin/streptomycin (Gibco) on 6-well plates coated with growth factor-reduced Matrigel (Corning) in an incubator at 37°C, 5% CO_2_, until 60-80% confluency was reached, at which point cells were split into new wells using ReLeSR passaging reagent (Stem Cell Technologies).

### PSC monolayer cardiac differentiation

Differentiation was performed using the small molecule Wnt modulation strategy as previously described^3^, referred to as monolayer 1 in the text), with small modifications. Briefly, differentiating cells were maintained in RPMI with B27 minus insulin from day 0-7 of differentiation and maintained in RPMI with B27 supplement (Thermo) from day 7-15 of differentiation. Cells were treated with 10 uM GSK inhibitor CHIR99021 (Selleck) for 24 hours at day 0 of differentiation and with 2 uM PORCN inhibitor, Wnt-C59 (Selleck), for 48 hours from day 3-5 of differentiation. The alternative differentiation protocol (referred to as monolayer 2) was described in Bertero et al, 2019^43^.

### Self-assembling heart organoid differentiation

Accutase (Innovative Cell Technologies) was used to dissociate PSCs for spheroid formation. After dissociation, cells were centrifuged at 300 g for 5 minutes and resuspended in Essential 8 Flex medium containing 2 µM ROCK inhibitor Thiazovivin (Millipore Sigma). hPSCs were then counted using a Moxi Cell Counter (Orflo Technologies) and seeded at 10,000 cells/well in round bottom ultra-low attachment 96-well plates (Costar) on day -2. The plate was then centrifuged at 100 g for 3 minutes and placed in an incubator at 37°C, 5% CO_2_. After 24 hours (day -1), 50 µl of media was carefully removed from each well and 200 µl of fresh Essential 8 Flex medium was added for a final volume of 250 µl/well. The plate was returned to the incubator for a further 24 hours. On day 0, 166µl (∼2/3 of total well volume) of media was removed from each well and 166 µl of RPMI 1640/B-27, minus insulin (Gibco) containing CHIR99021 (Selleck) was added at a final concentration of 4 µM/well along with BMP4 at 0.36 pM (1.25ng/ml) and ActA at 0.08 pM (1ng/ml) for 24 hours. On day 1, 166 µl of media was removed and replaced with fresh RPMI1640/B-27, minus insulin. On day 2, RPMI/B-27, minus insulin containing Wnt-C59 (Selleck) was added for a final concentration of 2 µM Wnt-C59 and the samples were incubated for 48 hours. The media was changed on day 4 and day 6. On day 6, media was changed to RPMI1640/B-27 (Gibco). On day 7, a second 4 µM CHIR99021 exposure was conducted for 1 hour in RPMI1640/B-27. Subsequently, media was changed every 48 hours until organoids were ready for analysis.

### Immunofluorescence

hHOs were transferred to microcentrifuge tubes (Eppendorf) using a cut 1000-μL pipette tip to avoid disruption to the organoids and fixed in 4% paraformaldehyde solution (dissolved in phosphate buffered saline (PBS)) for 30 minutes at room temperature. Fixationwasfollowedby3washesinPBS-Glycine(20mM)andincubationin blocking/permeabilization solution (10% Donkey Normal Serum, 0.5% Triton X-100, 0.5% bovine serum albumin (BSA) in PBS) on a thermal mixer (Thermo Scientific) at 300 RPM at 4°C overnight. hHOs were then washed 3 times in PBS and incubated with primary antibodies (see Table 1) in Antibody Solution (1% Donkey Normal Serum, 0.5% Triton X-100, 0.5% BSA in PBS) on a thermal mixer at 300 RPM at 4°C for 24 hours. Primary antibody exposure was followed by 3 washed in PBS and incubation with secondary antibodies (see Table 1) in Antibody Solution on a thermal mixer at 300 RPM at 4°C for 24 hours in the dark. The stained hHOs were washed 3 times in PBS before being mounted on glass microscope slides (Fisher Scientific) using Vectashield Vibrance Antifade Mounting Medium (Vector Laboratories). 90 µm Polybead Microspheres (Polyscience, Inc.) were placed between the slide and the coverslip (No. 1.5) to preserve some of the 3D structure of the organoids while accommodating the penetration capacity of the confocal microscope

### Confocal microscopy and image analysis

Samples were imaged using a confocal laser scanning microscope (Nikon Instruments A1 Confocal Laser Microscope) and images were analyzed using Fiji (https://imagej.net/Fiji). For tissue region quantification in the organoids, DAPI positive cells were used for normalization agains the target cell marker of interest across three z-planes throughout each organoid. A MaxEntropy threshold was used to measure the PECAM1+ tissue, discarding any stained region that took up less than 25 µm^2^ to remove noise.

### RNA sequencing and transcriptomic analysis

RNA was extracted at 11 different time points throughout the hHO fabrication and differentiation protocol shown in figure 1a. The time points are as follows: days 0, 1, 3, 5, 7, 9, 11, 13, 15, 17, and 19. At each time point, eight organoids were removed and stored in RNAlater (Qiagen) at -20°C until all samples were collected. RNA was extracted using the Qiagen RNEasy Mini Kit according to manufacturer instructions (Qiagen, 74104), and the amount of RNA was measured using a Qubit Fluorometer (ThermoFisher Scientific). RNA samples were sent to the MSU Genomics Core where the quality of the samples was tested using an Agilent 2100 Bioanalyzer followed by RNA sequencing using an Illumina HiSeq 4000 system. For RNA-seq sample processing a pipiline was created in Galaxy. Briefly, sample run quality was assessed with FASTQC, and alignment to hg38 was carried out using HISAT2. Counts were obtained using featureCounts and differential expression analysis was performed with EdgeR. Further downstream bioinformatic analysis were performed in Phantasus 1.5.1 (artyomovlab.wustl.edu/phantasus).

### Optical coherence tomography analysis

A customized spectral-domain OCT (SD-OCT) system was developed to acquire 3D images of the cardiac organoids. As shown in **Suppl. Fig. 5**, a superluminescent diode (SLD 1325, Thorlabs) was used as the light source to provide broadband illumination with a central wavelength of 1320nm and spectral range of 110 nm. The output of the SLD was split 50/50 with a fiber coupler and transmitted to the sample and reference arms, respectively. A galvanometer (GVSM002-EC/M, Thorlabs) was used to scan the optical beam in transverse directions on the sample. The SD-OCT setup used a custom-designed spectrometer consisting of a 1024-pixel line scan camera (SU1024-LDH2, Sensors Unlimited), an 1145 line pairs per mm diffraction grating (HD 1145 line pairs per mm at 1310 nm, Wasatch Photonics) and an f = 100 mm F-theta lens (FTH100-1064, Thorlabs). The sensitivity of the OCT system was measured as ∼104 dB when operating at 20 kHz A-scan rate. The axial resolution of the SD-OCT system was measured to be ∼7 μm in tissue. A 5X objective lens (5X Plan Apo NIR, Mitutoyo) was used to achieve a transverse image resolution of ∼7 μm, and the scanning range used for the cardiac organoids imaging was ∼2 mm X 2 mm. hHOs were placed into a 96-well plate with PBS, and imaged at 20-kHz A-scan rate. Obtained OCT datasets of the cardiac organoids were first processed to generate OCT images with corrected scales. Then OCT images were further de-noised using a speckle-modulation generative adversarial network^70^ to reduce the speckle noise. 3D renderings of OCT images were performed using Amira software (Thermo Fisher Scientific).

### Data Availability

All organoid data sets shown in this study are available at the National Center for Biotechnology Information Gene Expression Omnibus repository under accession number GSE153185. RNA-seq data from monolayer differentiation method 2 and fetal heart were obtained from GSE106690^43^.

### Statistical analyses

All analyses were performed using Excel or GraphPad software. All data presented a normal distribution. Statistical significance was evaluated with a standard unpaired Student t-test (2-tailed; P<0.05) when appropriate. For multiple-comparison analysis, 1-way ANOVA with the Tukey’s or Dunnett’ post-test correction was applied when appropriate (P<0.05). All data are presented as mean±SD and represent a minimum of 3 independent experiments with at least 3 technical replicates unless otherwise stated.

## Notes

### Competing Interest Statement

The authors have declared no competing interest.

### Summary of Updates

A problem with a missing figure in suppl. data has been fixed.

