## Supplementary data for "Generation of Heart Organoids Modeling Early Human Cardiac Development Under Defined Conditions"

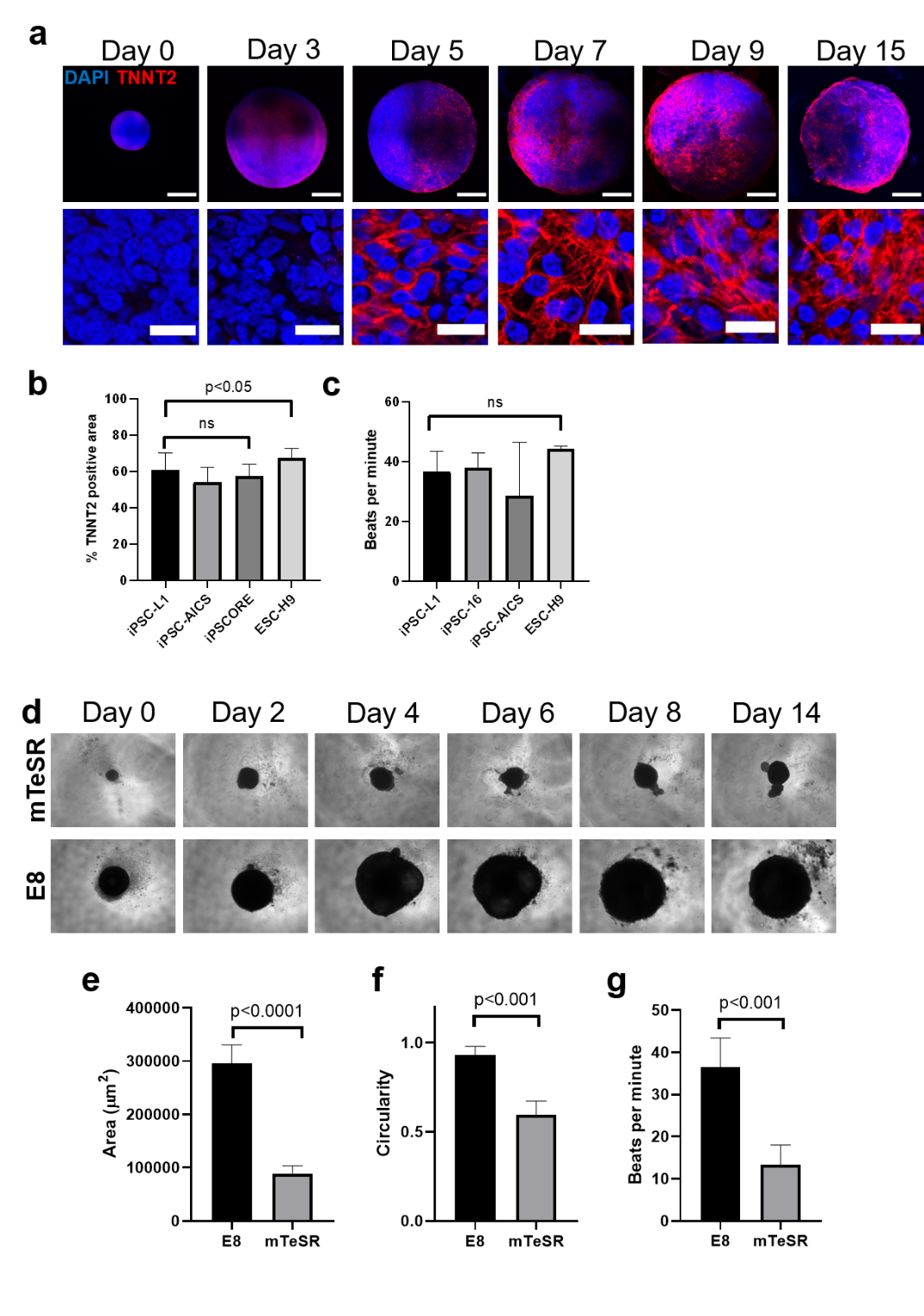

**Supplementary Figure 1. a,** Confocal immunofluorescent images for DAPI (blue) and TNNT2 (red), throughout hHO differentiation showing the development of sarcomeres; (Scale bar: top, 500µm; bottom 20 µm). **b,** Percentage of TNNT2+ area normalized to DAPI+ area in confocal images of hHOs, and **c,** beating frequency, in 3 iPSC lines and 1 ESC line. **d,** Light microscopy images of developing hHOs derived from iPSCs cultured in mTeSR Plus media (top) and Essential 8 media (bottom). **e,** Size of hHOs measured from light microscopy images, **f,** circularity of hHOs at day 14 and **g,** beat rate at day 14 for hHOs derived from iPSCs cultured in mTeSR Plus media and Essential 8 media; (Value = mean ± s.d., two-tailed, unpaired t-test).

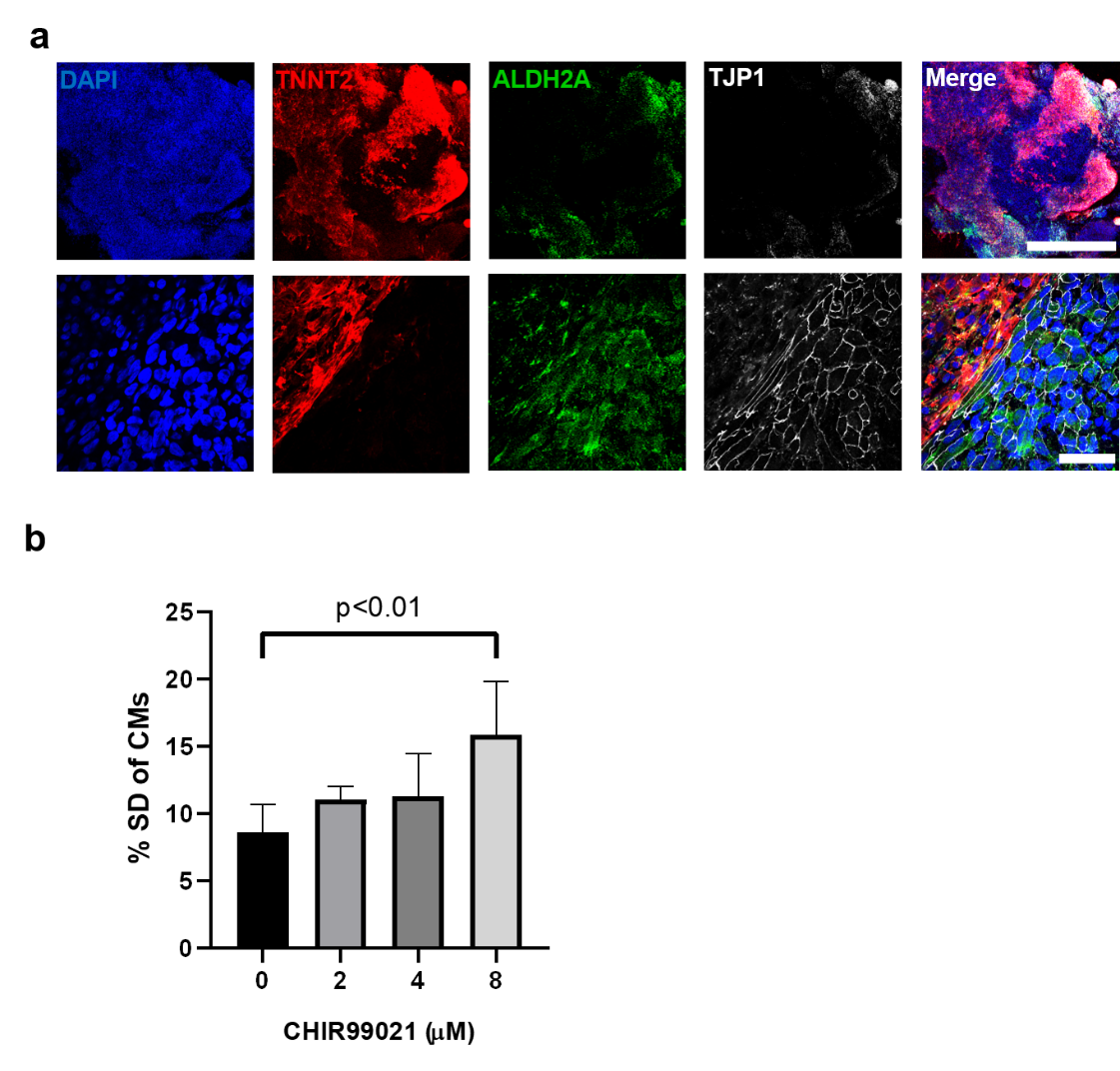

**Supplementary Figure 2. a,** Confocal immunofluorescent images for DAPI (blue) and TNNT2 (red), in hHOs showing epicardial markers ALDH2A (green) and TJP1 (white) near edge of the organoid. **b,** Percentage of standard deviations in TNNT2+ regions between hHOs treated with different concentrations of CHIR99021 at day 7; (Value = mean ± s.d, ordinary one-way ANOVA).

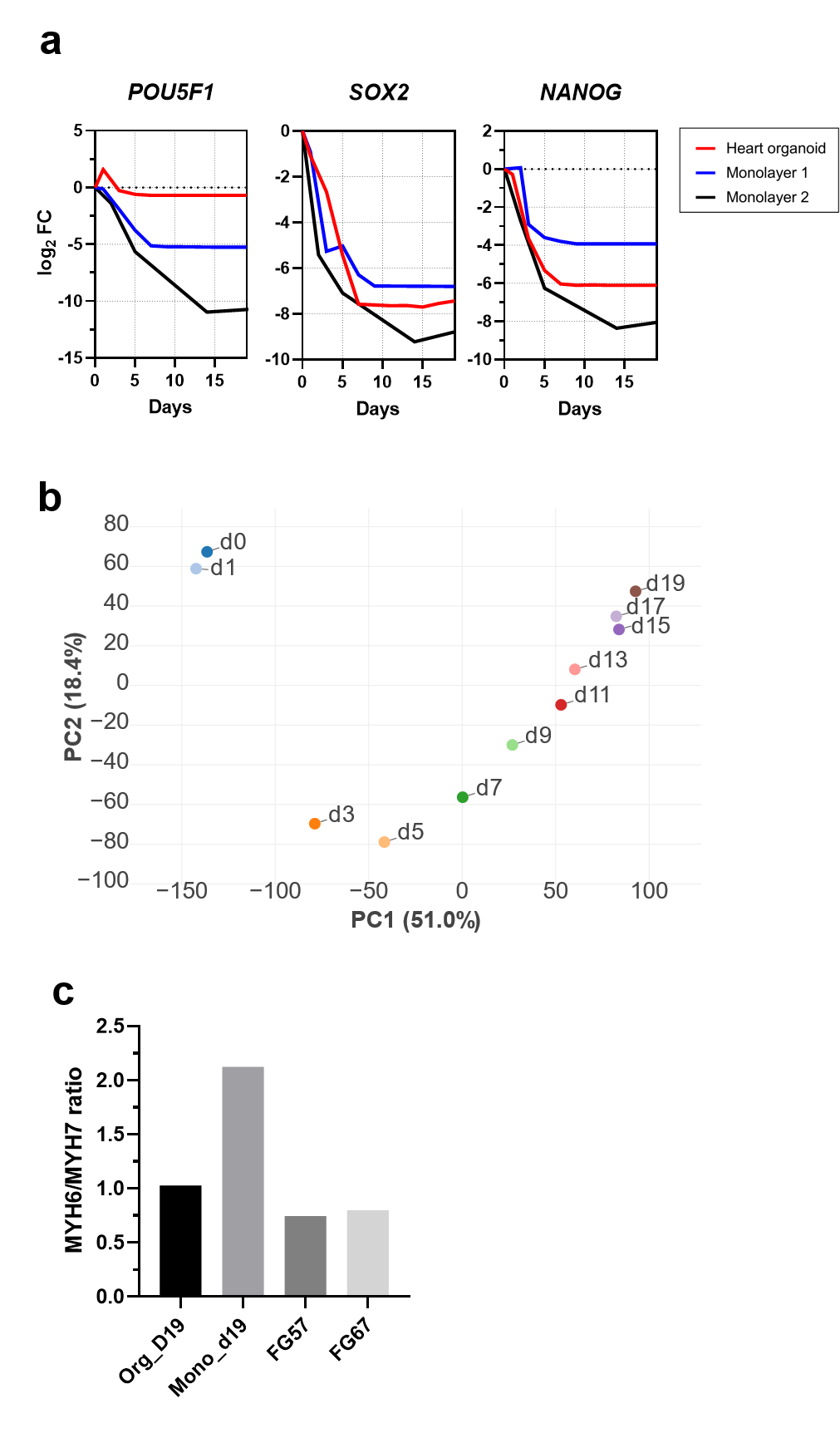

**Supplementary Figure 3. a,** Gene expression analysis indicating silencing of pluripotency network during heart organoid maturation (log_2_ fold-change vs. D0). Notice that POU5F1 expression is barely detectable in D0 organoids at the start of differentiation. **b**, Principal component analysis of heart organoid differentiation over time. **c**, The *MYH6/MYH7* ratio is a good indicator of cardiomyocyte maturation. Monolayer differentiation produces more mature and abundant cardiomyocytes. Heart organoid composition is closer to that of fetal hearts.

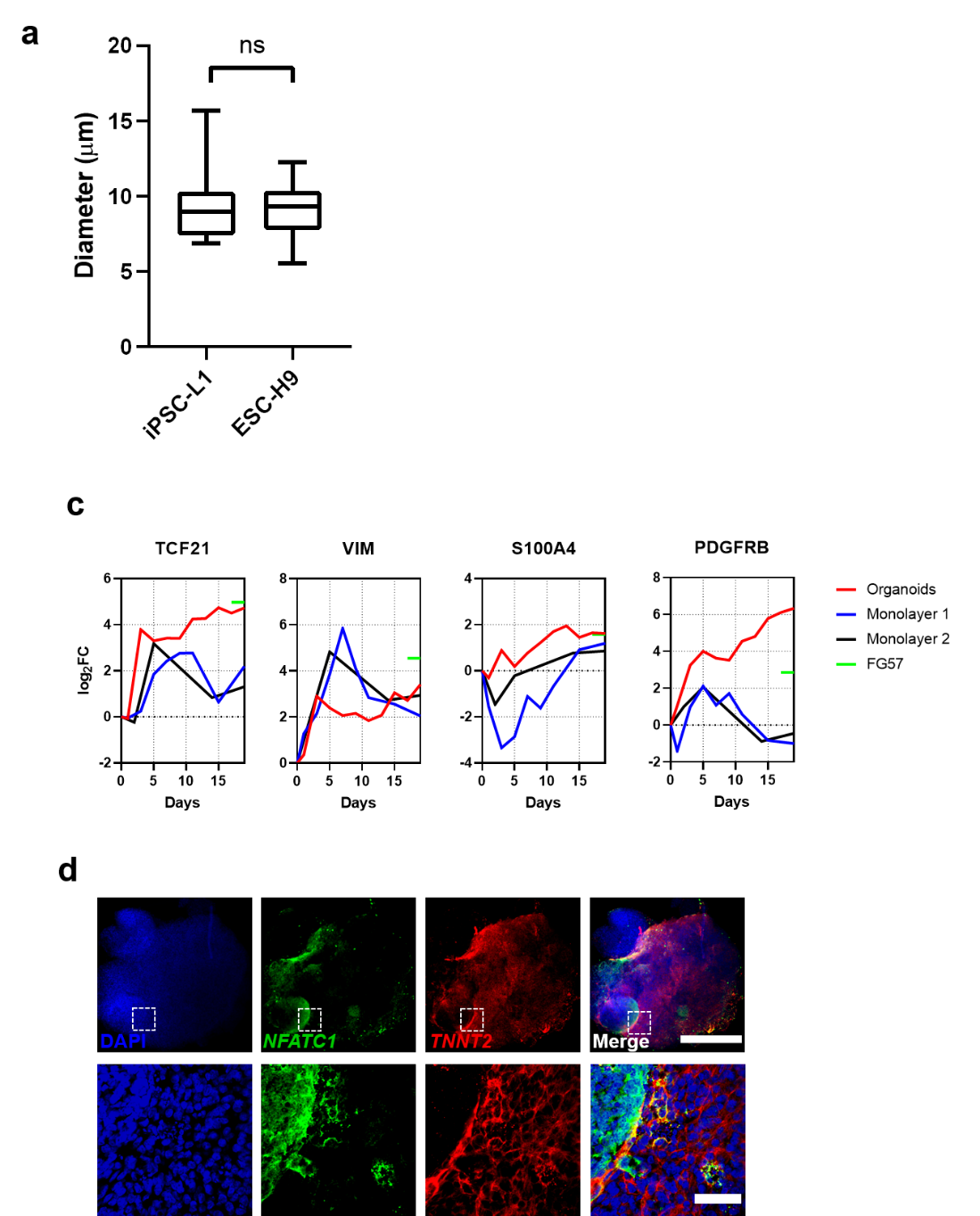

**Supplementary Figure 4. a,** Diameter of PECAM1+ vascular structures in iPSC-L1 and ESC-H9 (n=14); (Value = mean ± min., max., two-tailed, unpaired t-test). **b,** Gene expression analysis from RNA-Seq data of epicardial markers in hHOs (red) compared with monolayers (black and blue). Green line indicates expression level in fetal gestational day 57.

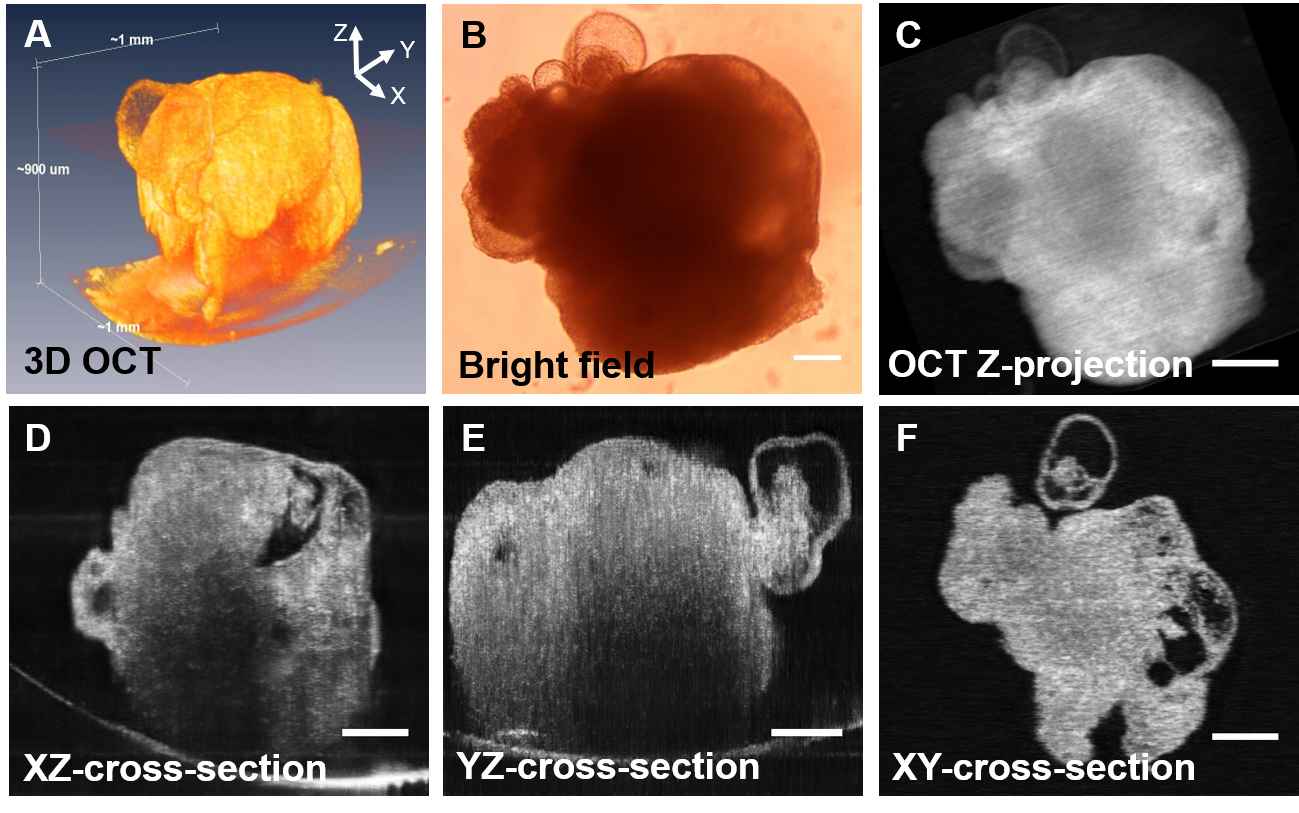

**Supplementary Figure 5.** OCT images and bright field image of human cardiac organoid. a) Three-dimensional (3D) rendering of an OCT image. b) Brightfield image of cardiac organoid, imaged with Olympus 4x objective lens. Scale bar: 200 µm. c) Projected *en face* OCT image. The image in is stacked along Z direction over 900 µm. d, e, f) 3D sectioning of OCT images in XZ, YZ and XY plane. Scale bars: 200 µm.

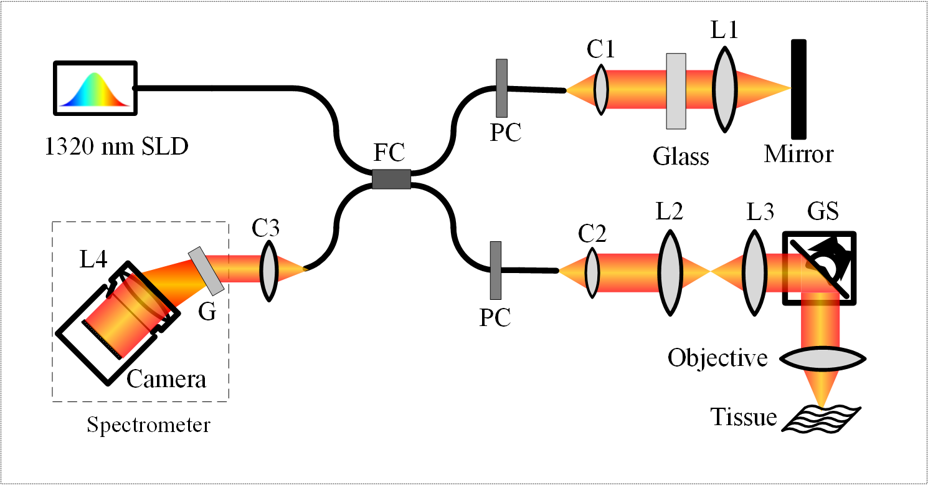

**Supplementary Figure 6.** Illustration of a custom SD-OCT imaging system, FC: fiber coupler, PC: polarization controller, C1-C3: collimator, L1 - L4: lens, GS: galvo scanner; G: grating.

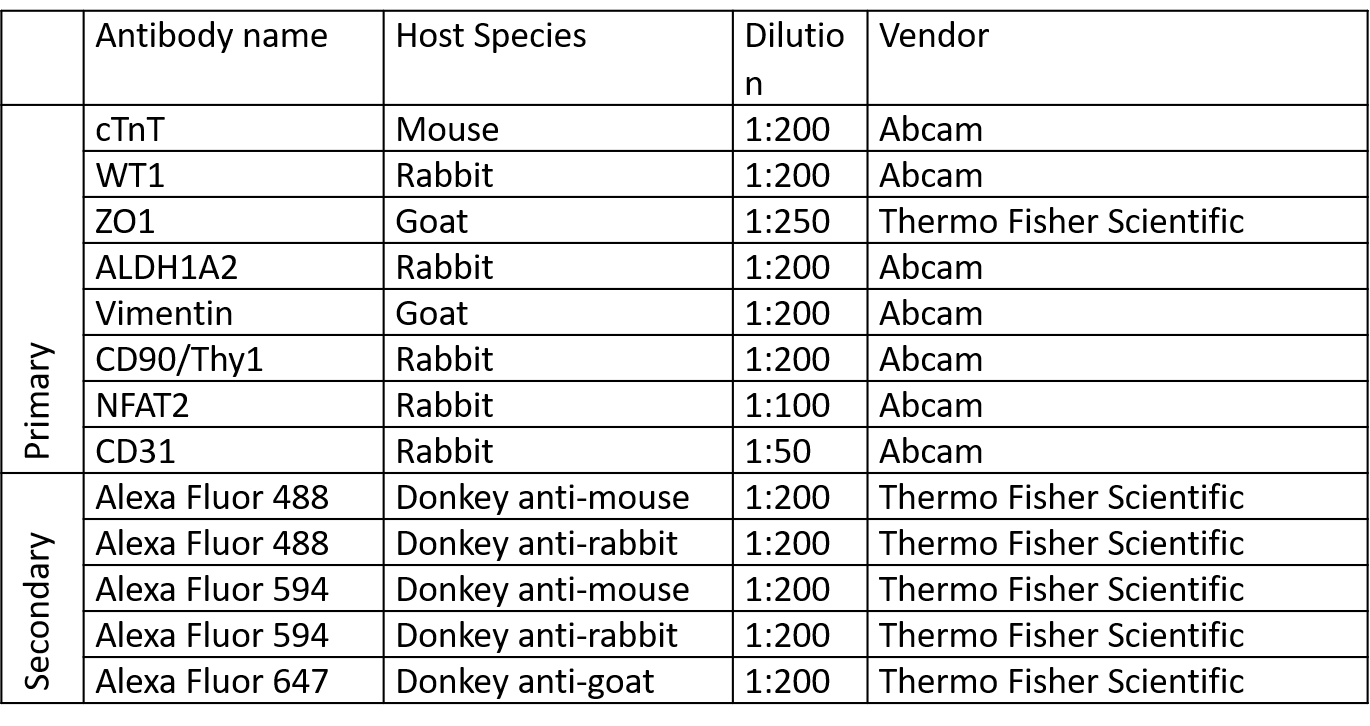

**Supplementary Table 1.** Antibodies used for immunofluorescence in this report.
